# Tracking global changes induced in the CD4 T cell receptor repertoire by immunisation with a complex antigen using local sequence features of CDR3 protein sequence

**DOI:** 10.1101/001883

**Authors:** Niclas Thomas, Katharine Best, Mattia Cinelli, Shlomit Reich-Zeliger, Hilah Gal, Eric Shifrut, Asaf Madi, Nir Friedman, John Shawe-Taylor, Benny Chain

**Affiliations:** UCL CoMPLEX, Physics Building, Gower Street, London, WC1E 6BT; UCL Division of Infection & Immunity, Gower Street, London, WC1 6BT; Weizmann Institute of Science, Rehovot, Israel, 7610001; UCL Computer Science, Malet Place, London, WC1E 6BT

## Abstract

The clonal theory of adaptive immunity proposes that immunological responses are encoded by increases in the frequency of lymphocytes carrying antigen-specific receptors. In this study, we measure the frequency of different TcRs in CD4+ T cell populations of mice immunized with a complex antigen, killed Mycobacterium tuberculosis, using high throughput parallel sequencing of the TcR beta chain. In order to track the changes induced by immunisation within this very heterogeneous repertoire, the sequence data were classified by counting the frequency of different clusters of short (3 or 4) continuous stretches of amino acids within the CDR3 repertoire of different mice. Both unsupervised (hierarchical clustering) and supervised (support vector machine) analysis of these different distributions of sequence clusters differentiated between immunised and unimmunised mice with 100% efficiency. The CD4+ T cell receptor repertoires of mice 5 and 14 days post immunisation were clearly different from that of unimmunised mice, but were not distinguishable from each other. However, the repertoires of mice 60 days post immunisation were distinct both from unimmunised mice, and the day 5/14 animals. Our results reinforce the remarkable diversity of the T cell receptor repertoire, resulting in many diverse private TcRs contributing to the T cell response even in genetically identical mice responding to the same antigen. Finally, specific motifs defined by short sequences of amino acids within the CDR3 region may have a major effect on TcR specificity. The results of this study provide new insights into the properties of the CD4+ adaptive T cell response.

## Introduction

Adaptive immunity is carried out by populations of B and T lymphocytes which collectively express a very large set of different antigen specific receptors created during haemopoesis by a unique process of somatic cell gene rearrangements. The clonal theory of immunity (Burnet, 1959) proposes that lymphocytes carrying receptors which specifically bind an antigen to which the immune system is exposed, for example during infection or vaccination, respond by proliferating and differentiating. This population of expanded and differentiated cells then confer on the system the ability to respond specifically to the antigen to which they had previously been exposed. The clonal theory therefore explains the immune system properties of specificity and memory. A prediction of this theory is that the frequency of lymphocytes which have been exposed to antigen (i.e. memory or effector cells) will be greater than the frequency of those which have not (i.e. naive). This prediction has been verified for T cells in a wide variety of models, using antigen specific readouts such as cytokine responses, and MHC multimer binding to identify expanded lymphocyte clones (Hataye et al., 2006; Moon et al., 2007; Catron et al., 2004). The selective expansion of specific clones has also been inferred from global measurements such as V region usage (Reuther et al., 2013) or spectratyping (a technique sometimes referred to as the immunoscope) (Russi et al., 2013). Previous studies have distinguished between private TcRs, found in one or a few individuals, and public TcRs found within the responding repertoire of a majority of individuals. The response to many antigens seems to consist of a mixture of public and private specificities (Cibotti et al., 1994; Menezes et al., 2007; Clute et al., 2010; Day et al., 2011).

The introduction of short read parallel high-throughput sequencing (HTS) provides an alternative approach to measuring lymphocyte receptor frequencies, allowing evaluation of the global receptor repertoire of particular lymphocyte populations. Rearranged receptor genes, or their mRNA products, are expanded and then sequenced directly, and the number of times each unique receptor sequence is found is simply counted. This approach can in principal generate an accurate estimate of the number of times each unique lymphocyte receptor is present in a particular population, and this information should reflect the prior antigen exposure of the individual. Several previous studies have already used HTS to reveal interesting properties of the BcR and TcR repertoire. Freeman et al. (2009) and Robins et al. (2009) used HTS to show non-uniform V(D)J gene segment usage in humans during recombination, which has been attributed to chromatin conformation (Ndifon et al., 2012). Complementary work on antibody repertoire diversity has also been conducted in the zebrafish again showing non-uniform V(D)J recombination that is qualitatively conserved between individuals (Weinstein et al., 2009). Weinstein et al. (2009) also show that this repertoire is shaped by maturity, with a greater skew in V(D)J usage observed at 2 months compared with 2-week old individuals. Other studies have used HTS to provide unexpected insight into the naive and memory T cell compartments, revealing that the memory compartment may be far more diverse than previously thought (Klarenbeek et al., 2010; Robins et al., 2009; Venturi et al., 2011). There have also been some more translational applications of HTS lymphocyte receptor sequencing, in the context of haemopoetic cancer diagnosis (Mamedov et al., 2011), bone marrow transplantation (Wu et al., 2012), and autoimmunity (Krell et al., 2013).

A major goal is to use the HTS lymphocyte receptor sequence data to identify antigen-specific changes in the repertoire at a global level. There remain some major challenges, however. Firstly, HTS generates primary sequence data, which can not easily be mapped onto three dimensional receptor conformation, much less onto intrinsic antigen specificity. Secondly, current technologies do not yet provide easy ways to link the two chains of the antigen specific receptor (heavy and light for antibody, alpha and beta or gamma and delta for T cells) at a single cell level. Indeed the majority of studies of T cell repertoires using HTS have focused on beta chains only. The antigen specificity of the receptor will depend on the pairing of a specific alpha and beta chain, and can therefore not be inferred from beta chains alone.

Despite these limitations, there are a number of indications that local features of protein primary structure may contain hidden information which reflects specific protein/protein interactions occurring at the level of fully folded tertiary or quaternary structure. One interesting example is the analysis of conserved amino acid pairs within a family of homologous proteins which has recently been used to predict with remarkable accuracy the structure of the fully folded protein on the basis of conserved protein/protein interactions (Hopf et al., 2012). Some success has also been achieved using primary protein structure to predict antibody/antigen docking (Brenke et al., 2012). From a machine learning perspective, these approaches are reminiscent of algorithms which use local low-level features such as individual words or image fragments to produce remarkably efficient classification of very complex large datasets, such as sets of documents or images. Surprisingly, good results can be achieved with little regard for the semantic content or meaning of these types of data. We thought that local sequence features could be used to define antigen-specific changes in the TcR repertoire following immunisation, reflecting underlying information on the nature of interactions between TcR and peptide-MHC complexes. In this study we develop an approach based on the well-studied bag-of-words (BOW) (Joachims, 1998; Csurka et al., 2004; Lowe, 1999) algorithm to categorise and classify sets of TcR sequences from immunised and unimmunised mice at different times post immunisation.

Our data highlight the extraordinary diversity of the T cell repertoire, which result in every individual mice expressing a majority of receptors which are unique to that individual. Within this ocean of diversity, conventional methods to identify the antigen specific component of the response by looking for shared expanded clones are problematic. Surprisingly, however, localised sequence features (similar short stretches of adjacent amino acids) can be used to generate a high dimensional feature space, in which the distinct experimental groups can be readily distinguished with a very high degree of accuracy. Short motifs in CDR3 primary sequence may therefore play an important role in determining TcR specificity. Our results suggest that the response to antigen may be an emergent property of the repertoire, dominated by clones found only in that individual (private specificities), and distributed over many lymphocytes each with different receptors. The response to antigen restricts the repertoire into a sub-space which is of lower dimension / reduced diversity compared with the full naive repertoire. In each individual, a number of different clones will be expanded out of the many clones within this sub-space. The analysis of short sequence features provides an approach to defining this restricted subspace of the repertoire based on the collective response of many individuals to the same challenge.

## Results

### Decombinator analysis of sequences from immunised and unimmunised mice

We collected spleens from unimmunised mice or mice immunised with heat-killed Mycobacterium tuberculosis 7, 14 and 60 days previously. CD4 T cells were isolated, and the TcRs sequenced as described in Methods. A summary of the analysis of the HTS data by using the modified Decombinator algorithm (methods and Supplementary Information) is given in Table 1.

**Table 1.** Summary of murine HTS TcR data used in this study. The analysis described in this study focused exclusively on functional TcRs. (1) The number of sequences generated which pass the sequence quality threshold. (2) The number of sequences recognized and classified by Decombinator as TcRs. (3) The number of sequences containing an in frame CDR3.(4) The number of TcRs obtained counting each distinct sequence once only. (5) The number of CDR3 amino acid sequences obtained, counting each CDR3 sequence once only.

|  | Sequences<sup>1</sup> | TcR sequences<sup>2</sup> | Functional TcRs<sup>3</sup> | Distinct nt sequences<sup>4</sup> | Distinct CDR3 sequences<sup>5</sup> |
| --- | --- | --- | --- | --- | --- |
| U | 1,788,551 | 428,326 | 404,032 | 186,599 | 113,429 |
| U | 3,430,471 | 873,607 | 831,605 | 303,266 | 171,004 |
| U | 1,023,799 | 172,105 | 164,114 | 97,601 | 69,125 |
| U | 2,394,189 | 175,924 | 166,398 | 74,141 | 48,244 |
| U | 2,949,666 | 347,621 | 326,310 | 106,112 | 65,806 |
| U | 3,840,471 | 216,790 | 207,135 | 38,902 | 27,660 |
| D5 | 10,005,718 | 2,034,715 | 1,931,868 | 720,436 | 415,436 |
| D5 | 2,770,773 | 930,044 | 884,619 | 408,830 | 256,116 |
| D5 | 4,479,250 | 1,589,692 | 1,512,244 | 432,490 | 267,451 |
| D5 | 2,480,475 | 834,436 | 795,269 | 205,329 | 138,134 |
| D5 | 10,413,014 | 2,730,565 | 2,576,696 | 1,008,680 | 558,64 |
| D5 | 10,578,666 | 2,050,542 | 1,946,156 | 531,214 | 317,637 |
| D14 | 5,689,589 | 1,884,138 | 1,792,945 | 570,080 | 346,164 |
| D14 | 4,297,876 | 1,320,231 | 1,254,062 | 517,549 | 323,291 |
| D14 | 6,415,997 | 1,260,362 | 1,205,562 | 371,107 | 236,863 |
| D14 | 4,251,640 | 1,258,960 | 1,203,263 | 372,241 | 236,572 |
| D14 | 4,220,098 | 842,190 | 799,261 | 318,764 | 206,532 |
| D14 | 5,829,487 | 1,182,640 | 1,131,125 | 413,942 | 260,594 |
| M2 | 4,497,142 | 424,287 | 406,912 | 154,001 | 107,337 |
| M2 | 6,570,454 | 441,799 | 422,609 | 111,140 | 82,213 |
| M2 | 6,754,251 | 764,024 | 732,688 | 153,383 | 110,533 |
| M2 | 5,303,993 | 357,340 | 341,430 | 110,816 | 83,392 |
| M2 | 3,601,458 | 339,273 | 321,471 | 203,614 | 139,381 |
| M2 | 7,124,801 | 410,596 | 387,232 | 177,126 | 121,348 |

In total, we analysed 120 million raw sequence reads, of which 19% were classified as specific TcRs by Decombinator. This proportion is similar to that observed using conventional pairwise alignment as described previously (Ndifon et al., 2012). The enormous diversity of the repertoire is emphasised by the fact that 76% of the identifiers were unique to a single mouse spleen. The proportion of unique, translated CDR3s was somewhat smaller (62%), reflecting the degeneracy of the genetic code. A few ‘public’ sequences were shared by all mice.

We first hypothesised that the repertoire of immunized mice might contain several identical expanded antigen-specific clones, and might therefore be more similar to each other than unimmunised mice. We estimated the similarity between mice using the Jaccard index, the ratio of the intersection to the union of the two samples. The distribution of Jaccard indices for all possible pairs of mice are shown in fig 2. Contrary to our prediction, this analysis did not demonstrate any greater similarity between pairs of immunised mice than between pairs of unimmunised mice. However, the Jaccard index for pairs composed of one immunised and one unimmunised mouse was significantly smaller than for pairs of two immunised, or two unimmunised mice (fig. 2, right). Immunisation therefore altered the repertoire state, but did not drive repertoire convergence.

Having established that we could detect a change in the CDR3 repertoire following immunisation, we investigated whether CDR3s from immunised mice shared primary protein sequence features which distinguished them from unimmunised mice. For this purpose we adapted the bag-of-words approach (also called the n-gram kernel) originally developed in the context of document recognition (Joachims, 1998), together with a clustering step to reduce the dimensionality of the vocabulary (see Methods for further details). The codebook used for classification was initially chosen to be one hundred clusters each containing a subset of contiguous, short (length *p*, where *p* typically = 3) stretches of amino acids, from the set of possible contiguous p-tuples. The similarity metric for clustering was based on individual amino acid Atchley factors (Atchley et al., 2005), reflecting similarities in physicochemical characteristics of the amino acids. For example, there are 8000 possible amino acid triplet combinations (20^3^), and these were allocated to 100 clusters, on the basis of similarity of Atchley factor scores. The contents of each cluster of amino acid triplets are given in Table S1 (SI), and the sizes of the 100 clusters are shown in Supp Fig 1.

**FIGURE 1:**
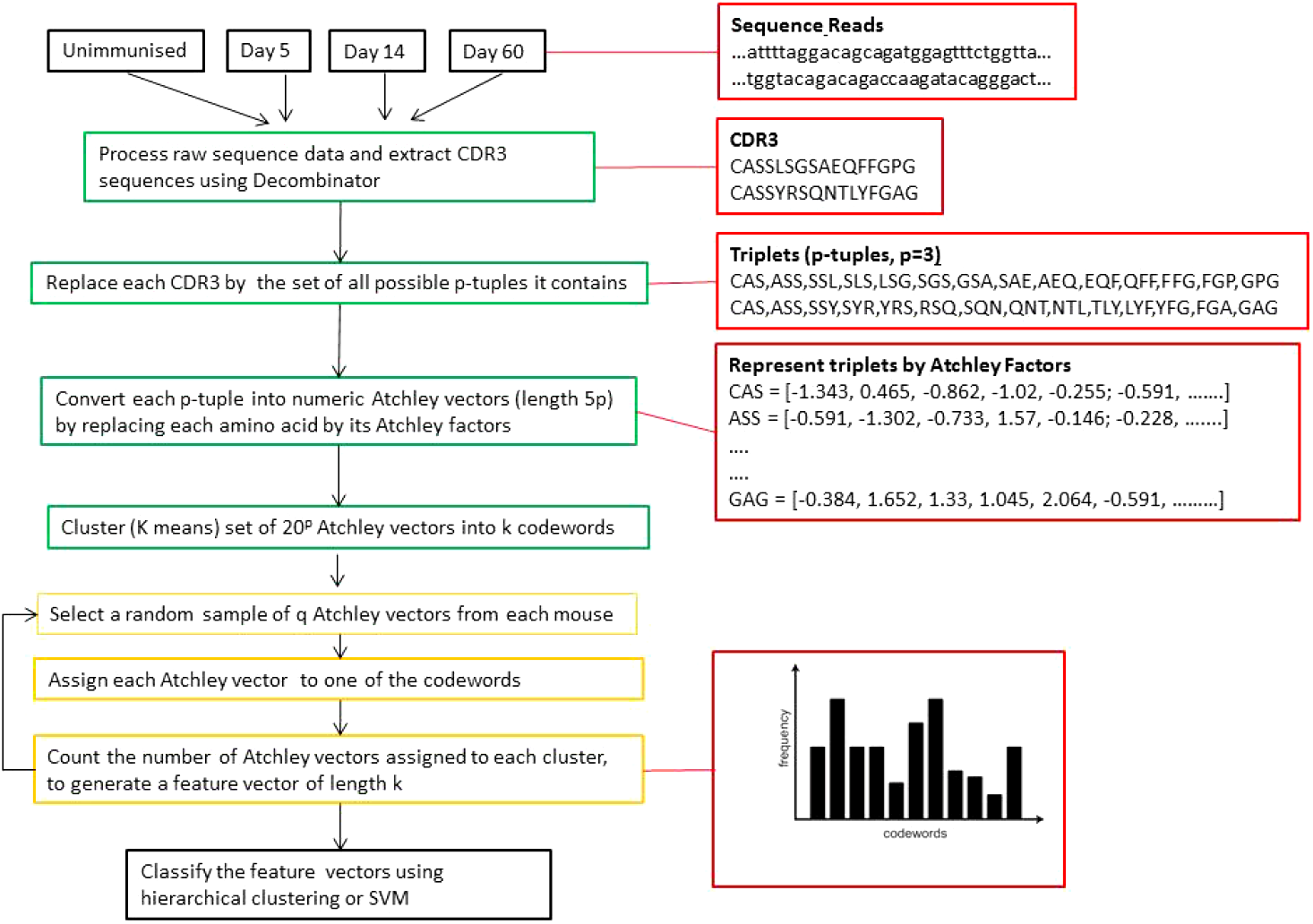
The computational pipeline for classifying TcR repertoires. A schematic of the computational pipeline is shown on the left, and a specific example for two arbitrary TcR beta sequences is shown on the right (with p = 3). CDR3 sequences are represented as a series of p-tuples (contiguous sequences of amino acids of length p). The p-tuples are then converted into numeric vectors of length 5p by representing each amino acid by its five Atchley factors. The set of all possible vectors is first clustered to build a codebook of *k* codewords via k-means clustering. A sample of q p-tuples from each mouse is then selected and mapped to the nearest codeword. Finally the number of p-tuples within each codeword for that mouse is counted. The sequence data from each mouse is therefore represented by a feature vector of length k, containing the frequency of each codeword within the sample. These k length vectors are then analysed by hierarchical clustering or support vector machines (SVM).

**FIGURE 2:**
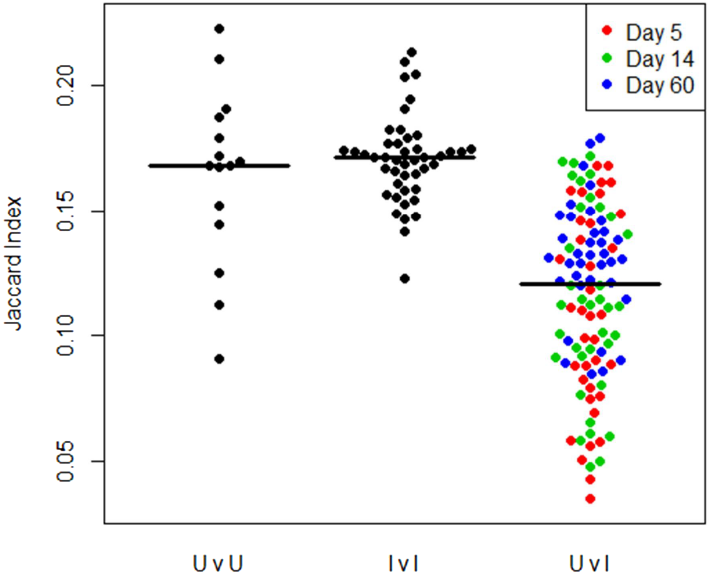
The similarity (Jaccard) index comparing all pairs of mice. Each dot represents the Jaccard index comparing all CDR3 sequences from two mice. CDR3 repertoires from pairs of unimmunized (U) mice, or pairs of immunised (I) mice, are significantly more similar (i.e. have a larger Jaccard index) than repertoires from pairs of mice where one mice is immunised, and one is not immunised. Horizontal black lines indicate mean of each population. (UvU versus UvI *p* = 6.9 × 10^−5^; IvI versus UvI, *p* < 2.2 × 10^−16^; UvU versus IvI *p* = 0.753).

Initially, we sampled 10,000 randomly selected amino acid triplets (i.e. p = 3) from the CDR3 region of each receptor, allocated each triplet to the nearest cluster, and then counted the total number of triplets within each cluster for that mouse. The set of primary sequences for each mouse were therefore mapped into a numeric feature vector of length 100. The results for all 24 mice are displayed in fig 3a, ordered by hierarchical clustering along both dimensions. Unsupervised clustering correctly separates all six unimmunised mice (on the left) from the immunised mice. Some additional structure is evident with most of the intermediate day 5/14 mice lying between the unimmunised and the 60 day group. Several different patterns of codeword distribution are observed. For example, codewords at the top of the heatmap become more highly represented in the repertoire of immunised mice, while codewords at the bottom are less represented.

**FIGURE 3:**
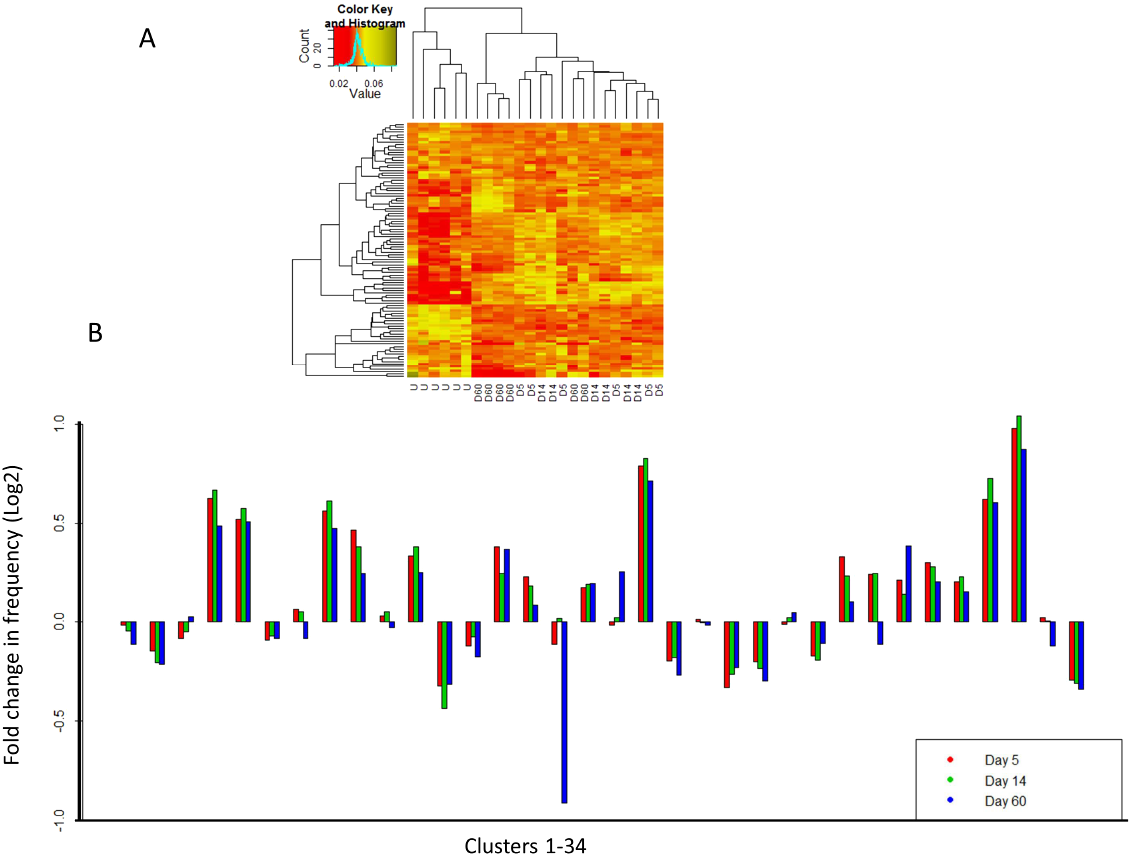
Codeword (clusters of triplets) distribution profiles differs between unimmunised and immunised mice. Each mouse was categorised as described in text, using k = 100, p = 3 (triplets), q = 10,000. a) The heatmap shows the relative proportion of sequences within each codeword (rows) for each mouse (colums). The data are clustered along both axes using euclidean distances and complete linkage method in the R function ‘hclust’. b) The data for each group of six mice are averaged, and shown relative to the unimmunised group. For clarity the data for only the first 34 codewords is shown.

A summary of the data, in which codeword cluster sizes from all mice in each group are averaged, and plotted as log ratio relative to unimmunised mice is shown in fig 3b. Consistent changes in cluster size are shown for many clusters, with a majority showing increases in the immunised groups. In most cases, the different timepoints behave in a similar manner, although large decreases in some clusters are seen specifically in the day 60 mice. The time dependence of the repertoire is examined further below.

In order to develop a classification tool with which we could explore the parameters of the bag-of-words algorithm, we constructed a multiclass SVM to classify the set of vectors obtained as above. Mice were classified as belonging to one of 4 classes, unimmunised/control, day 5, day 14 and day 60 post immunisation. Training and testing was performed using leave-one-out cross-validation on each mouse in turn.

The results of varying several of the parameters of the classification algorithms are shown in Table 2. Using a radial basis function for the SVM had little effect on classification efficiency, probably reflecting the inherent high dimensionality of the data. All further analysis was therefore carried out using linear SVM kernels. Decreasing the codebook size to 10 codewords compromised the success rate, as did decreasing the number of p-tuples sampled from 10000 to 1000. Increasing the sample size above 10000 had no further effect (not shown). Increasing p from 3 to 4 (i.e. quadruplets, rather than triplets) made little difference, although it was difficult to know if this was because efficiency was already close to 100%, or the additional information carried in the longer amino acid stretches was not informative. Interestingly, decreasing p to one (i.e. simply amino acid prevalence) retained some discriminative potential, albeit considerably reduced from *p* = 3 or 4.

**Table 2:**
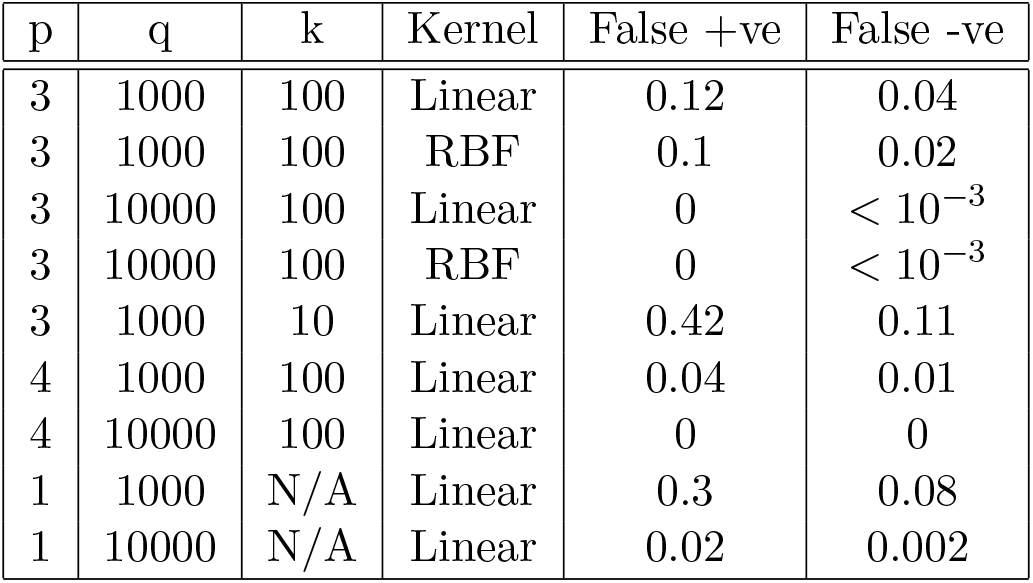
Efficient SVM classification using bag-of-words categorisation. Mice were classified as either immunised or unimmunised. False positive rate is the proportion of unimmunised mice classified as immunised. False negative rate is the proportion of immunised mice classified as unimmunised. k is number of codewords (clusters of Atchley vectors). q is number of p-tuples sampled from each mouse. p is the length of contiguous amino acid sequence. SVM uses either a linear or radial basis function (RBF) kernel.

We next examined classification efficiency retaining the separate time points post immunisation as distinct classes (fig. 4). For this purpose we used *k* = 100, *p* = 3, and *q* = 10000. Similar results were obtained for *p* = 4. As discussed previously, immunised and unimmunised mice are distinguished with 100% efficiency. As we expected, the repertoire of day 5 and 14 mice cannot be efficiently distinguished from each other, reflecting the fact that the T cell response at these two time points is likely to be similar. Interestingly, the repertoire of day 60 mice was often distinct from that of the earlier time points, although never returning to an unimmunized type. Thus the T cell repertoire appears to show a time-dependent change following immunisation. The efficiency with which day 60 and earlier time points could be distinguished varied between mice, suggesting that the time course of this evolution was somewhat variable between individuals.

**FIGURE 4:**
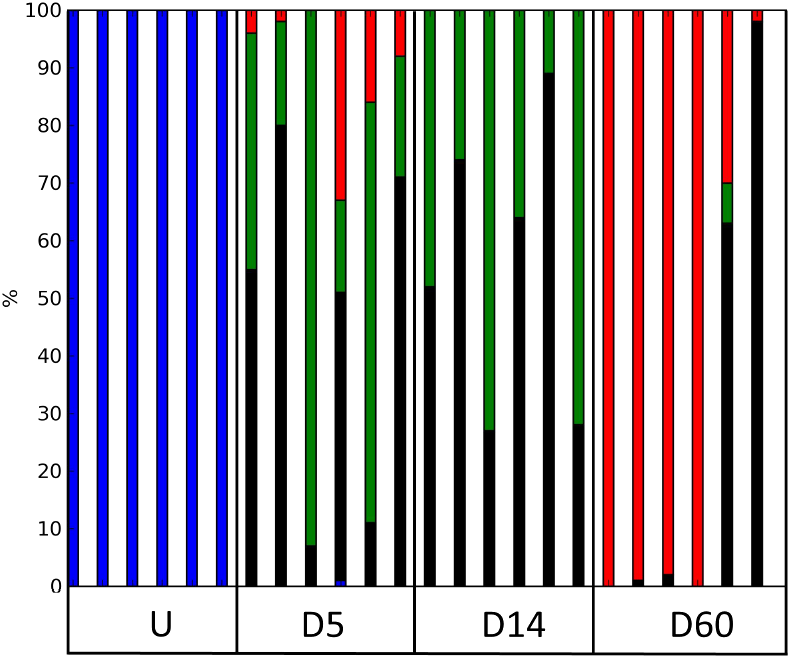
Time dependent changes in CDR3 repertoire following immunisation. The plot shows the distribution of classification results for each mouse using a leave-one-out linear SVM repeated 100 times. k = 3, p = 3, q = 10000. Blue: unimmunised; Black: day 5; Green: day 14; Red: day 60.

We wondered whether the distinctive features of the immunised repertoire were determined predominantly by a few very abundant T cells (i.e. highly expanded clones), or whether the repertoire was determined by many rare T cells sharing similar CDR3 sequence features. We therefore first ranked all the TcRs from each mouse according to the number of times it occurred in the sample, and then selected only CDR3 sequences from high frequency (top 10 percentile in clone frequency) or low frequency clones (bottom ten percentile) and repeated the analysis shown in fig. 4. Remarkably, equivalent classification efficiency was obtained using either high or low frequency clones. 100% correct classification was retained even further decreasing the cut off to the lowest 1% (i.e. almost all clones which appear only once in each sample). Increasing the cut off to use only those clones in the top 1 percentile (on average these appeared 40 times or more in a sample) increased the false positive rate to 16%. These results imply that the enrichment of specific sequence features in immunised mice reflect changes to many different T cell clones of both high and low frequency, rather than the dominant expansion of a few dominating antigen specific clones.

## Discussion

The computational pipeline presented above analyses the global T cell immune response to a complex antigen, killed Mycobacterium tuberculosis. This antigen contains many different proteins, and contains a large number of possible T cell epitopes. Despite this complexity, the results demonstrate a coherent but highly distributed set of responses, emerging from the background of the remarkable diversity and plasticity of the TcR generating system.

HTS of the T cell receptor repertoire of individual mice emphasised the size of the potential repertoire, consistent with previous reports (Ndifon et al., 2012). In order to simplify the computational aspects discussed in more detail below, we focus here exclusively on the CDR3 regions of the receptor, which are believed to contribute most to the interaction between TcR and the antigenic target peptide lying within the MHC groove (Garcia and Adams, 2005; Rudolph et al., 2006). The heterogeneity is highlighted by the observation that over 60% of the CDR3 repertoire is made up of unique sequences, and only a very small proportion are shared by all mice, even though all mice are derived from a well-established in-bred strain, and are therefore genetically very similar. On the basis of Jaccard index, this diversity extends equally to unimmunised and immunised mice. Thus immunisation, at least in this example, does not seem to result in the emergence of a large pool of shared identical CDR3 sequences. In contrast, the Jaccard index when comparing immunised and non-immunised mice is significantly lower than that obtained by comparing within either immunised or unimmunised groups. This suggests a model in which one heterogenous population of receptors changes as a result of immunisation to another equally heterogeneous, but nevertheless distinct, population. This picture of an immune response made up of frequency changes in many heterogeneous clones was confirmed by the further investigations detailed below.

Since we were unable to determine a clearly defined set of identical receptors which correlated with antigen response, we needed to devise a strategy to extract features which would reflect the similarities and differences between different data sets. This recognition had to be carried out in the absence of any direct information on potential antigen/receptor interaction for three reasons: the precise antigenic peptides processed and presented in the immunised mice were unknown, and because of the complexity of the antigen were probably diverse; the alpha chains pairing with individual beta chains were unknown; and the three dimensional conformation of the beta chains cannot be reliably inferred from the primary sequence. This problem shares features with the task of categorising texts or images, without recourse to high level (semantic or object-oriented) information about individual data sets. This type of problem has received an enormous amount of attention, and remarkable progress has been made over the past decades by using low level features, such as text strings or pixel clusters. As a first step, we adopted the simplest consecutive string kernel algorithm (often referred to as the bag-of-words method) (Lodhi et al., 2002; Joachims, 1998). In order to restrict the size of the feature space (there are 8000 possible triplet amino acid sequences, and 160000 quadruplets) we clustered the k-tuples, using Atchley factors (Atchley et al., 2005), which categorise amino acids according to a set of five orthogonal physical and chemical properties. Several alternative such classification schemes have been devised (e.g. Kawashima et al. (2008); Kidera et al. (1985)), and it will be of interest to see how these compare in the sequence classification type of problems investigated here.

Despite its simplicity, the feature space constructed from short consecutive amino acid p-tuples revealed a remarkably consistent time-dependent response to immunisation. Thus, while there was little sharing of identical sequences between groups of mice, shared patterns of sequences, defined by a particular distribution of p-tuples, was easily observed using both supervised and unsupervised classification methods. Although a few codewords (i.e. clusters of amino acid p-tuples) showed large differences between experimental groups, a substantial proportion of the codewords showed smaller but consistent changes. This suggested a large number of TcRs contribute to the antigen-driven changes in the composition of the repertoire, perhaps reflecting the complex nature of the antigen used in these studies. Indeed efficient recognition of different experimental groups required analysis of large numbers (q *≥* 10000) of codewords. Furthermore, the TcRs which defined the antigen specific repertoire were not confined to high frequency clones. Indeed, the classification efficiency of our SVM was unaffected by confining ourselves to clones which appeared only once in each sample. It should be noted that even ‘low frequency’ receptors may represent amplified clones, since sample size limits the lowest observable TcR frequency we can reliably see. Nevertheless, the data suggest a model where recognition of M. tb in these mice is distributed among many low and high frequency clones, sharing characteristic amino acid triples or quadruplets. At a molecular level, one might envisage that these selected subsequences may be directly interacting with specific features of antigenic processed peptides exposed at the surface of the MHC binding groove (Garcia and Adams, 2005). In fact a number of distinct TcRs are likely to interact with a single peptide/MHC complex, with a spectrum of different affinities (Birnbaum et al., 2012). In such a model, although the overall recognition between TcR and MHC/peptide is mediated at the level of tertiary or quaternary structure, and therefore not reducible to linear sequence features, the interaction between CDR3 and a specific peptide/MHC may impose constraints which are observable at the level of short contiguous amino acid sequences. Similar constraints have been demonstrated to characterise conserved protein/protein interactions (Hopf et al., 2012) in large evolutionary related protein families.

The majority of previous studies have measured individual T cell antigen specific responses without reference to TcR sequence (e.g. using MHC multimer binding or cytokine responses). Previous analysis of specific virus specific CD8 cells have described both private and public (shared) clones between individuals (Klarenbeek et al., 2012; Dash et al., 2011). More global approaches to detecting and quantifying receptor diversity have used spectratyping to obtain a profile of CDR3 lengths, or flow cytometry to quantify V region usage (Ciupe et al., 2013; Pannetier et al., 1995; Faint et al., 1999; Lim et al., 2002). Both techniques have given interesting insights into clonal expansions associated with a variety of antigen-driven responses, although the sensitivity limits the detection of small clones. Spectratyping can be extended to give sequencing data, but this is a laborious and low throughput process. We predict that, as larger sequence data sets become available from HTS approaches, the extent of diversity in the antigen-driven TcR repertoire response will increase dramatically. The present study is confined to a single antigen, in a single inbred strain of mice. Additional studies are in progress to extend the data sets to better defined model antigen systems, for example focusing on one individual MHC/peptide response. Preliminary results suggest the response to such weaker and narrower antigen stimuli are more subtle, and will require more sophisticated analysis. Many extensions of the current approach are possible. For example, the feature space can be extended, by including V and J region information, positional information in the context of the p-tuple within the CDR3, and the inclusion of non-continuous string kernels which are now computationally feasible (Lodhi et al., 2002).

The results described above offer an intriguing insight into the nature of an immune response. On the one hand, the success of classification methods using fairly simple low level features of protein sequence offer hopeful indications for applying this sort of approach to analysis of clinical samples for the prognosis, diagnosis or stratification of patients in the context of both infectious and non-infectious (e.g. cancer, autoimmunity, transplantation) disease. On the other hand, if further studies generalise our observation of a ‘distributed’ immune response, in which a response is carried by large numbers of different low frequency clones with shared features, this will pose some formidable computational challenges. Robust experimental pipelines, improved HTS technology and application of the latest advances in machine learning will all be required, but such combinations are likely to provide new insights into the function of the adaptive immune system, and ultimately translational benefits in the clinical context.

## Methods

### T Cell Receptor Library Construction

Briefly, C57BL/6 mice were immunised with freeze-dried Mycobacterium tuberculosis H37RA in a water/oil emulsion (Complete Freunds Adjuvant, Difco). Groups of six mice were sacrificed at day 5, day 14 and day 60, and spleens were harvested, CD4 T cells were purified by magnetic bead sorting and processed for mRNA extraction. Ovalbumin (100 ug/mouse) was included in some immunisations, but the presence or absence of ovalbumin had no effect on the analysis reported below and is not considered further in this study. Six further mice were left unimmunised. Subsequently, mRNA from the isolated T cells was reverse transcribed, amplified and sequenced as described in detail in (Ndifon et al., 2012). Briefly, reverse transcription was carried out using a *Cβ*-specific primer linked to an *Illumina* 3’ sequencing adapter. The resulting cDNA product was amplified with a multiplex PCR using a set of 23 *V β*-specific primers. Each *V β*-specific primer was anchored to a restriction site sequence for a restriction enzyme (AcuI) that was used to cleave part of the primer sequence, to ensure good coverage of the hypervariable CDR3 with a single short Illumina read. This was followed by ligation of a Illumina 5’ adapter, which was linked to a 3-bp barcode sequence at its 3’ end, and a second round of PCR amplification using primers for the 5’ and 3’ Illumina adapters. Final PCR products were gel purified and sequenced using the Genome Analyzer II.

### Low level processing of Illumina sequence reads to generate protein CDR3 sequences

Sequences obtained using this protocol were 55 base pairs long, spanning the highly diverse CDR3 region. Following the methodology described in (Thomas et al., 2013), we represent each distinct TcR sequence read in terms of its constituent V and J gene segments, the number of V and J germline nucleotide deletions and the string of nucleotides found between the VJ junction, including any remnants of the D gene segment. Thus, this approach classifies each TcR sequence in terms of 5 variables, mitigates for sequencing error within V or J regions and determines the correct reading frame to extract the translated CDR3 region.

The short length of the sequences made direct use of Decombinator (Thomas et al., 2013) problematic for unambiguous assignment of V and J gene segments, as the optimal unique tags that recognise the distinct V and J gene segments are located outside the sequenced window, and are necessarily located far enough from the 3’ and 5’ ends of the V and J gene segments respectively to ensure the deletion of nucleotides from the gene segment ends do not affect their detection. Additionally, the V*β* region located between the primer and the VD junction is very similar across all 23 mouse V*β* genes, making creation and detection of unique tags difficult, and the variability in the length of the CDR3 region means the number of J gene nucleotides that are present in each read varies from significantly, making selection of a single identifying J keyword difficult. Therefore, some modifications to the Decombinator (Thomas et al., 2013) pipeline were introduced which are described in detail in *Supplementary Information*.

### Atchley Factors

Amino acids were grouped according to similar physicochemical properties, using Atchley factors, which provide a sequence metric (Atchley et al., 2005) representing each amino acid by a vector that accounts for five orthogonal characteristics, based on polarity, secondary structure, molecular volume, codon diversity and electrostatic charge. Thus, TcR protein sequences are represented by a series of 5 dimensional vectors encoding functional properties of individual amino acids.

### The Bag-of-Words Approach

The basic strategy, used successfully in text, image and also protein sequence classification, is to define a large set of low level features (codewords) within the data, which is variously referred to as a codebook, dictionary or vocabulary. These can be individual words of text, image features or any other simple descriptive features (see e.g (Joachims, 1998; Csurka et al., 2004; Lowe, 1999)). Individual data items are then defined by how frequently each codeword of the vocabulary is found within that specific piece of data. Each data item is therefore converted into an k-dimensional vector, where k is the size of the vocabulary. Finally the overall data is classified into one of two or more sets, using one of a number of high dimensional classification tools. The pipeline is illustrated in fig. 1.

In the specific example examined here, the vocabulary is initially defined as all possible sets of contiguous, short (length *p*, where *p* typically = 3) stretches of amino acids (called p-tuples) within the set of CDR3 regions. These represent the features within the data. The p-tuples are then converted into a numeric Atchley vector of length 5*p*, by representing each amino acid by its five Atchley factors, which describe various of its physico-chemical properties. In order to reduce the size of the vocabulary to manageable size *k* (typically 100 ‘codewords’), the set of observed vectors is clustered. A sample of Atchley vectors is first generated from a set of sequences selected randomly from all experimental groups. This set is clustered into k clusters using k-means clustering. These k clusters represent the codebook (Supplementary data, fig S1 and Table S1). Once the codebook is defined, the repertoire of sequences from each mouse can be mapped to this codebook. A new set of sequences is selected from each mouse. The CDR3s from this set are converted into p-tuples, and then into Atchley vectors of length 5*p* as described above. Each Atchley vector is allocated to the nearest cluster. Once all vectors are allocated, the number of vectors within each cluster is counted, and converted into a proportion of the total number of Atchley vectors selected (q). In this way, each repertoire is mapped into a single k-dimensional vector.

These *k*-dimensional feature vectors are then classified using either unsupervised (hierarchical clustering) or supervised learning algorithms. For the latter we focused on support vector machines (SVM), which seek a linear hyperplane that separates observations from two (or more) distinct classes. The separating hyperplane is found such that the *margin* between the hyperplane and the nearest observations from the training data from each class is maximised, and the observations that define the size of the margin are termed support vectors, lending the method its name.

In cases where the data are not linearly separable, soft-margin optimisation is carried out via the introduction of slack variables (Cortes and Vapnik, 1995) to determine the optimal hyperplane for non-linearly separable data. The data can also be transformed into a higher dimensional space, where linear separation may be possible, using the so-called kernel trick (reviewed in Cristianini and Shawe-Taylor (2000)).

SVM was performed by using the e1071 package in R, and models are trained and tested using leave-one-out cross-validation. Multi-class discrimination is carried out internally in e1071 using a ‘one-against-one’ model.

